# A stagewise response to mitochondrial dysfunction in mitochondrial DNA maintenance disorders

**DOI:** 10.1101/2023.10.24.563545

**Authors:** Amy E. Vincent, Chun Chen, Tiago Bernadino Gomes, Valeria Di Leo, Tuomas Laalo, Kamil Pabis, Rodrick Capaldi, Mike Mauresch, David McDonald, Andrew Filby, Andrew Fuller, Diana Lehmann Urban, Stephan Zeirz, Marcus Deschauer, Doug Turnbull, Amy K Reeve, Conor Lawless

## Abstract

Mitochondrial DNA deletions clonally expand in skeletal muscle of patients with mtDNA maintenance disorders, impairing mitochondrial oxidative phosphorylation dysfunction. Previously we have shown that these mtDNA deletions originally arise and accumulate in the perinuclear mitochondria causing localised mitochondrial dysfunction before spreading through the muscle fibre. We believe that mito-nuclear signalling is a key contributor in this process.

To further understand the role of mito-nuclear signalling, we use imaging mass cytometry to characterise the levels of mitochondrial respiratory complexes I-IV and ATP synthase alongside a mitochondrial mass marker, in a cohort of patients with mtDNA maintenance dosirders. We then expanded this panel to include protein markers of key signalling pathways to investigate the cellular response in fibres with different combinations of oxidative phosphorylation dysfunction and in ragged red fibres.

We find CI and CIV deficiency to be most common, with a smaller proportion of cells that are also CIII and/or CV deficient. Interestingly, we also note that in cells deficient for one or more complexes, any complexes which are not deficient are commonly upregulated beyond the increase of mitochondrial mass typically observed in ragged red fibres. We further find that oxidative phosphorylation deficient fibres exhibit an increase in abundance of proteins involved in the proteostasis e.g. HSP60 and LONP1, and mitochondrial protein synthesis e.g. PHB1. Our analysis suggests that the cellular response to mitochondrial dysfunction changes depending on the combination of deficient oxidative phosphorylation complexes in each cell.

## Introduction

Mitochondrial DNA (mtDNA) maintenance disorders arise due to mutations in genes responsible for the maintenance and replication of mtDNA (e.g. POLG, TWNK, RRM2B) ^1–4^. Patients either have childhood-onset mtDNA depletion syndromes e.g. Alpers’ syndrome or adult-onset multiple mtDNA deletions disorders, e.g., Chronic Progressive External Ophthalmoplegia (CPEO). In the latter case, multiple mtDNA deletions form throughout life independently in individual cells post-mitotic cells, such as skeletal muscle fibres. Some of these deleted genomes clonally expand with age, causing mitochondrial dysfunction and progressive mitochondrial myopathy.

Why these mtDNA deletions persist and accumulate and whether they have a role in the pathogenesis remain insufficiently understood. Is the process purely stochastic or is there a driver behind the accumulation? Furthermore, how do the cells adapt and respond to the increasing levels of mtDNA deletions? Low levels of mtDNA deletions can persist in cells without an effect on mitochondrial function. However, as the mtDNA deletion level increases, it surpasses a threshold level, above which oxidative phosphorylation (OXPHOS) deficiency in the cell occurs^5^. Furthermore, it has previously been demonstrated in patients with single, large-scale mtDNA deletion patients that the threshold level of deletion required for complex I (CI) and complex IV (CIV) deficiency differs and is dependent on the location of the mtDNA deletion in the mtDNA^6^. As such, some mtDNA deletions will lead to a simultaneous deficiency in both CI and CIV, whilst others will lead to deficiency in one complex and then the other. These findings in single, large-scale mtDNA deletions, and further work in multiple mtDNA deletions^7^, support the idea that in a cell with multiple mtDNA deletion species, the location of the mtDNA deletion and the level of each mtDNA deletion will impact upon the overall OXPHOS dysfunction for that cell.

Muscle biopsies from patients with mutations in the mtDNA present upon sequential cytochrome oxidase (COX) succinate dehydrogenase (SDH) histology with a mosaic pattern of normal and deficient fibres ^8^. In addition, a proportion of deficient cells often develop into ragged red fibres (RRFs)^9^. RRFs are a pathological hallmark of mitochondrial myopathy and appear to be the product of disrupted mitophagy ^10^ and potentially also increased mitochondrial biogenesis in response to mitochondrial dysfunction. The development and progression of mitochondrial dysfunction at a single muscle fibre level appears to be stage- wise, but less is known about how the cellular response evolves through these stages of disease.

Metabolic analysis of patients with multiple mtDNA deletions has demonstrated activation of the mitochondrial integrated stress response (ISRmt), through mammalian target of rapamycin 1 (mTORC1) signalling in skeletal muscle^11^. This signalling process is thought to drive the progression of mitochondrial disease, with mTORC1 especially activated in RRFs ^10^. Indeed, rapamycin decreased the amount of affected cells and mtDNA deletions even in advanced disease ^11^. Previous work in C. elegans has also implicated the activation the retrograde mito-nuclear signalling which is thought to; 1) initiate a metabolic shift towards glycolysis ^12,13^ and 2) upregulate mitochondrial biogenesis allowing the clonal expansion of mtDNA deletions ^14,15^. Recent work has demonstrated that mitophagy levels exhibit a mosaic pattern in skeletal muscle and that mitophagy occurs mainly in the perinuclear region^10^. Conversely in RRFs, mitophagy was found to be stalled with an abundance of lysosomes observed across the muscle fibre^10^. This is likely to contribute towards the increase of mitochondrial mass in RRFs, and pharmacological induction of mitophagy was found to partially rescue this disease phenotype ^10^.

Much of the previous work on the cellular response to mitochondrial dysfunction assesses tissue homogenates, which does not allow comparisons between normal and OXPHOS deficient cells. In comparison, work from Murgia, et al. ^16^ compared a small number of cytochrome c oxidase (COX) positive and negative muscle fibres using proteomics. They found two clusters of proteins to be enriched in COX negative fibres: 1) pyruvate dehydrogenases (PDH) complex and the tricarboxylic acid (TCA) cycle and 2) chaperones. This second group included PDH1 and 2 and proteases such as Lon Peptidase 1 (LONP1). However, this analysis was done on a small number of cells and only looked at CIV (COX) deficiency.

We have previously hypothesised that clonal expansion of mtDNA deletions initially occurs in the perinuclear region based on the observation that the smallest regions of mitochondrial OXPHOS deficiency are found in the perinuclear area^17^. It was therefore suggested that the proximity to the nuclei facilitates mito-nuclear retrograde signalling and upregulation of mitochondrial biogenesis. We further observed that retrograde signalling factor G Protein Pathway Surppressor 2 (GPS2)^18^ and the chaperone protein Heat Shock Protein 60 (HSP60), were upregulated in perinuclear foci of deficiency, as were the markers of mitochondrial biogenesis peroxisome proliferator-activated receptor gamma coactivate 1-alpha (PGC1α) and Transcription factor A, mitochondrial (TFAM)^17^. However, these proteins were not upregulated in all foci of deficiency and the use of immunofluorescence prohibited the simultaneous investigation of greater than five targets and the ability to correlatate the analysis of a greater number of proteins. Imaging mass cytometry (IMC) allows the simultaneous analysis of a larger number of protein targets in a single tissue section whilst still maintaining enough spatial resolution for single and sub-cellular analysis^19^.

To more fully characterise the mitochondrial OXPHOS deficiency and understand the progressive response of cells to OXPHOS deficiency in single muscle from patients with mtDNA maintenance disorders, we harness the power of IMC and previously developed methods^20,21^, to simultaneously probe 19 signalling proteins, four mitochondrial markers, and a muscle cell marker. In doing so, we are able to look at not just the single cell levels but also the subcellular levels and distribution of a range of cell signalling proteins and understand how these change as mitochondrial dysfunction progresses.

## Methods

### Patients and tissue collection

Muscle biopsies of patients (n=12) with genetically and clinically characterised mitochondrial disease due to nuclear mutations were included in the study. Patients had mtDNA maintenance disorders with genetically confirmed and segregating pathogenic variants in POLG, TWNK and RRM2B. Clinical data and genotypes are summarised in Table 1. Genetic and data analysis were performed in compliance with protocols approved by the Ethical Committee of the Martin Luther University Halle-Wittenberg. Written informed consent was obtained from all participants prior to study inclusion. Three of the cases (P07, P08 and P16) were investigated with informed consent by the Newcastle and North Tyneside Local Research Ethics Committees (REC ref. 2002/205). Control tissue was acquired with prior informed consent from people undergoing anterior cruciate ligament surgery, following approval by Newcastle and North Tyneside Local Research Ethics Committees (REC ref. 12/NE/0395).

**Table 1.** Information and molecular diagnosis of patients with mtDNA maintenance disorders.

|  | Skeletal muscle histochemistry | Genotype | Previously reported |
| --- | --- | --- | --- |
| P1 | 30-40% COX-ve; 3-5% RRF | Dominant p.(Phe961Ser) **POLG** | P1 in Lehmann et al (2019) |
| P2 | 10% COX-ve; 2% RRF | Dominant p.(Lys319Glu) **TWNK** | P3 in Lehmann et al (2019) |
| P3 | 8-10% COX-ve; 1% RRF | Dominant p.(Met455Thr) **TWNK** | P4 in Lehmann et al (2019) |
| P4 | slightly increased number of COX-ve; 1% RRF | Dominant p.(Arg354Pro) **TWNK** | P5 in Lehmann et al (2019) |
| P5 | 9% COX-ve; increased number of RRF | p.(Tyr955Cys) **POLG** | P2 in Lehmann et al (2019) |
| P6 | 22% COX-ve; 8% RRF | Recessive p.(Arg186Gly) and p.(Thr218Ile) **RRM2B** | P8 in Lehmann et al (2019) |
| P7 | 20% COX-ve; 10% RRF | Recessive p.(Gly848Ser) and p.(Ala899Thr) **POLG** | P9 in Lehmann et al (2019) |
| P8 | 22% COX-ve; 1% RRF; | Recessive homozygous p.(Trp748Ser) **POLG** | P10 in Lehmann et al (2019) |
| P9 | 10% COX-ve; 1% RRF | Recessive p.(Arg467Thr) and p.(Thr919Leu) **POLG** | P13 in Lehmann et al (2019) |
| P10 | 17%, COX-ve, 9% RRF | Recessive homozygous p.(Gly426Ser) **POLG** | P14 in Lehmann et al (2019) |
| P11 | n.k | Recessive p.Ala467Thr; p.Thr251Ile/p.Pro587Leu **POLG** |  |
| P12 | n.k | Recessive p.Ala467Thr; p.Ala467Thr **POLG** |  |

### Antibody selection and panel design

The antibodies chosen were a combination of previously optimised mitochondrial and signalling targets^17,20,22^ Fmaz, with additional new targets from cell signalling markers. Targets were selected to cover key pathways of interest which included ISRmt, mitophagy, mitochondrial biogenesis, mitochondrial retrograde signalling, glycolysis and mitochondrial proteostasis (Table 2). To be included antibodies had to be available in a protein free format (except for GPS2, which we purified using an Abcam Protein G purification kit). All antibodies selected were known to be specific, either based on available data from the literature, the manufacturer or from our own testing (knock down experiments)^22^. All antibodies were optimised for the best working concentration by testing a range of dilutions using immunofluorescence. The highest possible dilution that produced a signal to noise ratio greater than 50 arbitrary fluorescence units and the expected staining pattern was selected. Subsequently, staining of sections with each antibody in combination with a mitochondrial mass or OXPHOS marker, was used to rank the chosen targets by signal intensity and to assign antibodies to appropriate metals, looking to minimise possible cross- talk as described previously ^20^.

**Table 2.** Antibodies used in the IMC experiments.

| Antibody | RRID | Description | Product code/Supplier | Metal Tag | Experiment | Final dilution |
| --- | --- | --- | --- | --- | --- | --- |
| anti-ATF4 | AB_2616025 | Transcription factor cAMP response elements | 11815/Cell Signalling | <sup>161</sup>Dy | 2 | 1 in 50 |
| anti-ATF6 | AB_2799696 | Transcription factor cAMP response elements | 65880/Cell Signalling | <sup>164</sup>Dy | 2 | 1 in 50 |
| anti-ATP5B | AB_301438 | Complex V | ab14730/Abcam |  | 1 | 1 in 50 |
| anti-BCL1 | AB_2704709 | Autophagy | StressMarq | <sup>169</sup>Tm | 2 | 1 in 50 |
| anti-CLPP | AB_1078538 | Protease/UPRmt | HPA010649/Sigma | <sup>166</sup>Er | 2 | 1 in 100 |
| anti-CHOP | AB_2089254 | Protease/UPRmt | 2895/Cell Signalling | <sup>167</sup>Er | 2 | 1 in 50 |
| anti-COX4+4L2 | AB_10862101 | Complex IV | Ab110261/Abcam |  | 1 | 1 in 50 |
| anti-MTCYB | * | Complex III | * | <sup>115</sup>In | 1 and 2 | 1 in 50 |
| anti-DJ1 | AB_11179085 | Protein repair | 5933/Cell Signalling | <sup>149</sup>Sm | 2 | 1 in 50 |
| anti-DRP1 | AB_11178938 | Density-regulated protein | 5391BF/Cell Signalling | <sup>168</sup>Er | 2 | 1 in 50 |
| anti-Dystrophin | AB_2091355 | Muscle membrane marker | Mab1645/Millipore | <sup>176</sup>Yb | 1 and 2 | 1 in 50 |
| anti-GPS2 | * | Retrograde stress signalling | * | <sup>154</sup>Sm | 2 | 1 in 50 |
| anti-HKII | AB_2232946 | Metabolism, Akt and TORC signalling | 2867/Cell Signalling | <sup>142</sup>Nd | 2 | 1 in 50 |
| anti-HSP60 | AB_399008 | Chaperone protein | 611562/BD Transduction systems | <sup>152</sup>Sm | 2 | 1 in 50 |
| anti-HTRA2 | AB_2280094 | Serine protease Htra2, mitochondrial | AF1458/R&D Systems | <sup>165</sup>Ho | 2 | 1 in 100 |
| anti-LONP1 | AB_1079695 | Lon protease homolog, mitochondrial | HAP002192/Sigma | <sup>172</sup>Yb | 2 | 1 in 100 |
| anti-MTCOI | AB_2084810 | Complex IV | ab218215/Abcam | <sup>174</sup>Yb | 1 and 2 | 1 in 50 |
| anti-MTHFD2 |  | One carbon cycle | ab151447/Abcam | <sup>153</sup>Eu | 2 | 1 in 50 |
| anti-MTND4 | * | Complex I | * | <sup>164</sup>Dy | 1 | 1 in 50 |
| anti-NDUFB8 | AB_10859122 | Complex I | ab110242/Abcam | <sup>160</sup>Gd | 1 and 2 | 1 in 50 |
| anti-OSCP | AB_10887942 | Complex V | ab110276/Abcam | <sup>161</sup>Dy | 1 | 1 in 50 |
| anti-PDH-E2 | Ab_10858998 | Glycolysis | ab110332/Abcam | <sup>150</sup>Nd | 2 | 1 in 50 |
| anti-PGC1α | AB_2268462 | Master regulator of mitochondrial biogenesis | AB3243/Millipore | <sup>174</sup>Yb | 2 | 1 in 100 |
| anti-PHB1 | AB_823689 | Proteostasis | 2426/Cell Signalling | <sup>171</sup>Yb | 2 | 1 in 50 |
| anti-SDHA | AB_301433 | Complex II | ab15715/Abcam | <sup>153</sup>Eu | 1 | 1 in 50 |
| anti-SIRT3 | AB_10861832 | NAD-dependent protein deacetylase sirtuin-3, mitochondrial | ab86671/Abcam | <sup>159</sup>Tb | 2 | 1 in 50 |
| anti-TFAM | AB_10900340 | Transcription Factor A Mitochondrial | ab119684/Abcam | <sup>156</sup>Gd | 2 | 1 in 50 |
| anti-UqCRC2 | AB_2213640 | Complex III | ab14745/Abcam | <sup>174</sup>Yb | 1 | 1 in 50 |
| anti-VDAC1 | AB_443084 | Voltage-dependent anion-selective channel protein 1. Mitochondrial mass marker. | ab218214/Abcam | <sup>147</sup>Sm | 1 and 2 | 1 in 50 |

### Antibody conjugation

Antibodies were conjugated to heavy metals using the protocol and Maxpar™ antibody labelling kit from Fluidigm, as described previously ^20^.

### Imaging mass cytometry

Cryosections of muscle tissue at 6μm were collected onto glass slides in arrays of four tissue sections per slide to reduce the number of slide changes needed and the amount of user interaction required during tissue ablation. Sections were labelled with the panel of antibodies listed in Table 2, as described previously ^20^. Two IMC experiments were performed, one where the sections were labelled with a full panel of mitochondrial targets and another where four key mitochondrial targets (MTCO1, NDUFB8, VDAC and MTCYB), were used alongside a range of cell signalling markers. Table 2 includes an experiment number to describe inclusion into panel 1 or 2. Regions of interest were selected to maximise the number of single fibres collected per case, whilst aiming to collect a single ROI per section to allow for spatial analysis using x and y co-ordinates.

### Image segmentation in Mitocyto

Images were exported as single channel 8-bit TIFFs from MCD viewer software. Mitocyto an in-house segmentation tool for skeletal muscle was used to automatically segment muscle fibres based on the Dystrophin (a cell membrane marker) signal, as reported previously ^20^. Longitudinal fibres, those with freezing artefact or tissue folding were removed by manual correction. Following image analysis, mean protein expression levels and morphological measurements taken for each fibre from each subject were exported and merged into a .csv file for data analysis.

### Single pixel segmentation

TIFFs were imported into a project in QuPath v0.4.0^23^. Segmentation of individual muscle fibres was performed manually based on the cell membrane (Dystrophin signal), VDAC (mitochondrial mass) and nuclear labelling using Iridium intercaltor. Fibres that were longitudinal or exhibited freezing artefact were excluded from analysis. Care was taken to include the myonuclei which were identified within the myofibril as denoted by the dystrophin cell membrane marker. Each cell was divided up into individual pixels (1μm^2^ as captured by the Hyperion) and mean intensity per pixel extracted.

### Data analysis

#### Definition of respiratory chain deficiency

All data analysis was performed using R v2021.09.1^24^. OXPHOS deficiency was defined by one of two methods. Firstly, to assess mitochondrial deficiency across all five OXPHOS complexes we used the tool PlotIMC as previously reported to allow interactive analysis of the comprehensive dataset ^20^. However, when it came to defining respiratory chain deficiency for the assessment of signalling changes we employed unsupervised Gaussian Mixture Modelling (GMM) of mitochondrial proteins using the MClust package ^25^. We used GMM to cluster fibres into two groups based on the OXPHOS complex marker (NDUFB8, MTCYB or MTCO1) and mitochondrial mass marker (VDAC), to define deficiency of CI, CIII and CIV respectively. These clusters were then compared to control data to classify which cluster represented “like control” fibres and which cluster represented OXPHOS deficient or “not-like controls”. The advantage of GMM is that it allowed us to first split fibres within a single patient without relying on the control samples and then to compare these two fibre groups to the control data to determine which group was control-like and therefore could represent the internal control fibres for each sample. We found that this approach led to proportions of deficient fibres that were closer to what we would expect based on prior investigation with techniques such as COX/SDH.

Visual examination of 2Dmito plots (Figure S1) of MTCO1, NDUFB8 and MTCYB against VDAC1 were used to assess the effectiveness of GMM for clustering fibres as OXPHOS normal or deficient. GMM accurately identified deficient cells for MTCO1 and NDUFB8 (Figure S1A and B). However, GMM was was not always successful in identifying a cluster that repsented fibres deficient for MTCYB (Figure S1C), due to the upregulation of MTCYB in fibres that are not deficient but are deficient for either NDUFB8 and MTCO1. Therefore the “not like control” group generally contained cells that had both lower or deficient levels compared to controls and the with higher levels compared to controls. Finally, RRFs were manually identified by systematic analysis of the IMC images, based on high VDAC1 levels and muscle fibre morphology (Figure S1D).

#### Definition of nuclear and cytoplasmic pixels

Single pixel data for each segmented cell was extracted using QuPath. By thresholding the Iridium intercalator signal (DNA1) across all pixels each pixel was defined as either nuclear or cytoplasmic. A range of threshold values were tested and all pixels in a section plotted and colour coded for nuclear or cytoplasmic location, these were then compared with the IMC pseudoimages to validate the approach. Based on this spatial validation, a cut off of DNA1>15 was used. After identifying nuclear and cytoplasmic pixels a mean nuclear signal and mean cytoplasmic signal was calculated for each protein in each individual cell.

#### Bayesian Estimation

Bayesian Estimation modelling^26^ was performed to compare cells with different combinations of mitochondrial OXPHOS deficiency in individual patients and to compare RRFs to controls, using the BEST package in R.

#### R script availability

All R scripts and pipelines used for analysis are available on GitHub (link).

## Results

### Characterisation of mitochondrial oxidative phosphorylation deficiency

Using IMC, we were able to simultaneously assess markers for OXPHOS complexes I-V and a mitochondrial mass marker VDAC1 ^20,21^, allowing to more thoroughly characterise mitochondrial OXPHOS deficiency in a cohort of patients with genetically ad clinically characterised mtDNA maintenance disorders (Figure 1A). As previously reported by Warren et al., we used plotIMC, an online interactive platform for dynamic visualisation of IMC data (http://mito.ncl.ac.uk/vincent_2019/). PlotIMC has three graphical representations of the data: i) 2D mitoplots showing each marker plotted on the Y axis against VDAC1 on the X axis (Figure S2A), ii) stripcharts of mean intensity for all channels (Figure S2B) and iii) stripcharts of theta (the angel formed by a line between each point in the 2D mitoplot and the origin and the X-axis, Figure S2C). One of the most interesting observations from looking at the data on PlotIMC is that the only patient with a large number of fibres with intensities lower than controls in all complexes is P11. For most patients the lowest intensities are either comparable to controls or less than 30 fibres have intensities lower. The real deficiencies are only noticeable when we look at the theta stripcharts, where mitochondrial mass (VDAC1) has been accounted for. This suggests that most cells with a deficiency in OXPHOS complexes also have an increase in mitochondrial mass.

**Figure 1.**
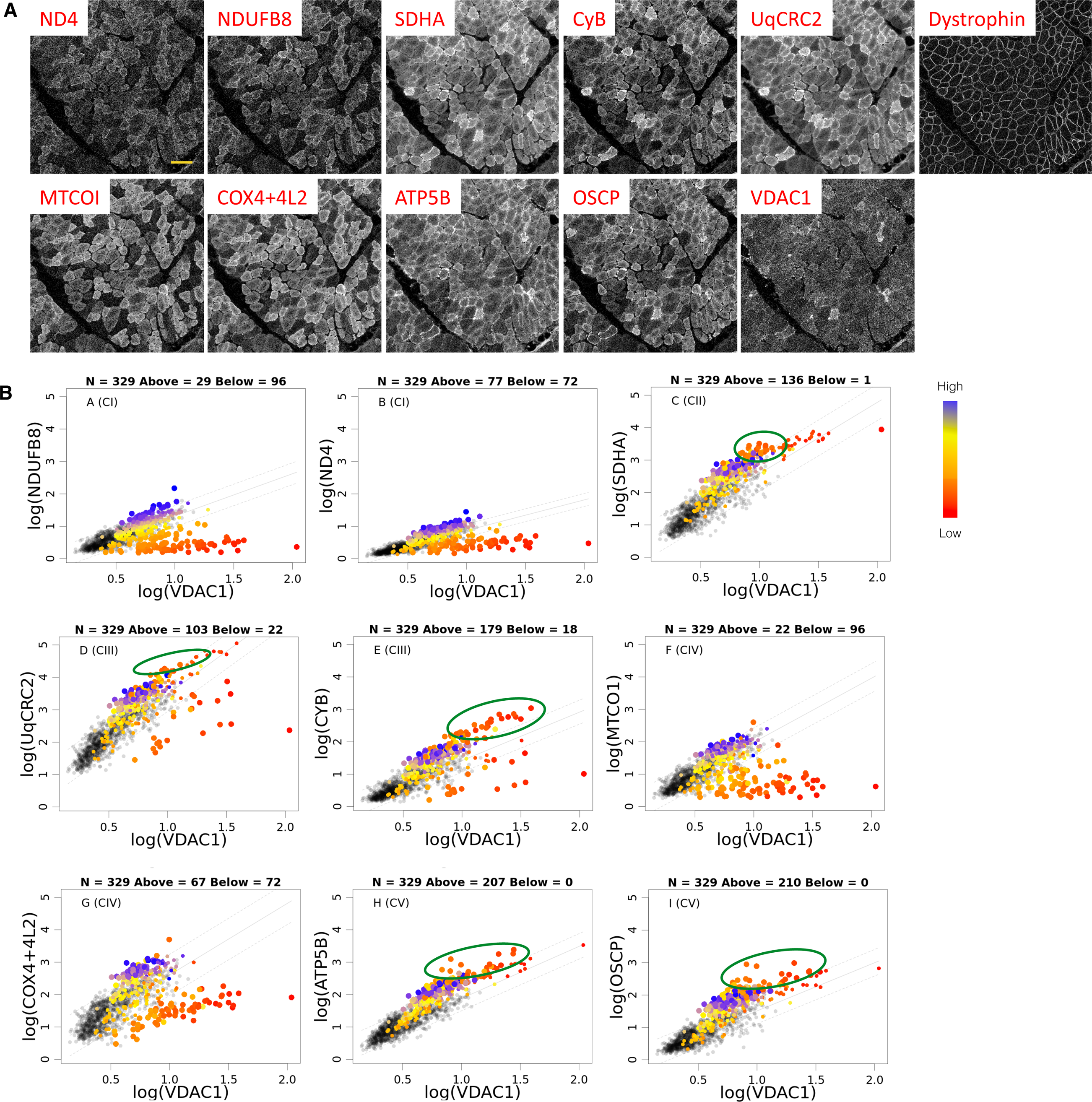
Quantification of subunits of oxidative phosphorylation complexes. (A) Example of imaging mass cytometry pseudo-images to assess representative subunits of oxidative phosphorylation (OXPHOS) complexes (ND4, NDUFB8, SDHA, CyB, UqCRC2, MTCOI, COX4+4L2, ATP5B, OSCP), mitochondrial mass (VDAC1) and a muscle fibre membrane marker (Dystrophin). (B) 2D mitoplots for each OXPHOS subunit plotted against the mitochondrial mass VDAC1. Each point represents a single muscle fibre. Points representing patient fibres are colour-coded based on the levels of the complex I (CI) subunit NDUFB8 (blue being high and red low). Grey points represent control fibres. Dashed lines represent the 95% predictive interval of the control data. Above each plot are quoted the total number of fibres (N), and the numbers of fibres classified as being above or below the upper and lower limits of the 95% predictive interval, respectively. Green circles are used to indicate fibres with increased protein level due to CI deficiency.

We observed cells with a range of combinations of OXPHOS deficiency profiles including isolated CI, III, or IV deficiency and combined deficiencies including CI with either CIII, CIV or both, as well as combined deficiency of CI, CIII, CIV and CV (Table 3). Such combinations of deficiency and the frequency at which they arise fit with the previously reported spectra of mtDNA deletions^7^. The deletions reported previously by Lehmann, et al.^7^. show deletions impacting the region from MT-ND6 to MT-ND1, fitting with CI and CIV deficiency being most frequent followed by CV and then CIII. The percentage of these deficiencies in patients bore no obvious relationship with the underlying nuclear gene mutated (Table 1), similar to that previously reported^7^. As with previous work by Warren, et al. ^20^ looking at a range of mtDNA mutations, we observed an upregulation of complexes that did not exhibit defiency at a single cell level, including CII (SDHA), CIII (MTCYB), CV (ATP5B and OSCP) (indicated by the green circles Figure 1B). We also observed strong correlations between subunits of the same complex, with high R^2^ values (Spearman’s rank correlation) observed for subunits from the same complex in all patients (Figure S3); MTND4 and NDUFB8 (R^2^=0.91-1), UqCRC2 and MTCYB (R^2^=0.79-0.99), MTCO1 and COX4+4L2 (R^2^= 0.92-1) and ATP5B and OSCP (R^2^=0.85-1), with the exception of CI in P11 and CIII in P12. Each of the OXPHOS markers are highly correlated with each other in controls, however these correlations often fall apart in patients, for, example in P02 NDUFB8 vs OSCP has R^2^ = 0.0.

**Table 3.** Summary of the proportion of fibres classified as belonging to each oxidative phosphorylation deficiency combination.

|  | CV | CIV | CI | CIII | CI+CIII | CI+CIV | CI+CV | CIII+CIV | CIII+CV | CIV+CV | CI+CIII+CIV | CI+CIV+CV | CIII+CIV+CV | All | None |
| --- | --- | --- | --- | --- | --- | --- | --- | --- | --- | --- | --- | --- | --- | --- | --- |
| P01 | 0.000 | 0.027 | 0.023 | 0.000 | 0.004 | 0.000 | 0.000 | 0.000 | 0.000 | 0.000 | 0.025 | 0.000 | 0.000 | 0.000 | 0.779 |
| P02 | 0.000 | 0.043 | 0.043 | 0.000 | 0.000 | 0.000 | 0.000 | 0.000 | 0.000 | 0.000 | 0.055 | 0.000 | 0.000 | 0.000 | 0.666 |
| P03 | 0.012 | 0.018 | 0.005 | 0.002 | 0.001 | 0.000 | 0.000 | 0.000 | 0.000 | 0.012 | 0.018 | 0.001 | 0.000 | 0.001 | 0.926 |
| P04 | 0.022 | 0.040 | 0.045 | 0.000 | 0.003 | 0.000 | 0.006 | 0.002 | 0.000 | 0.022 | 0.040 | 0.040 | 0.001 | 0.030 | 0.682 |
| P06 | 0.009 | 0.019 | 0.041 | 0.000 | 0.009 | 0.000 | 0.000 | 0.000 | 0.000 | 0.009 | 0.073 | 0.001 | 0.000 | 0.001 | 0.710 |
| P07 | 0.010 | 0.025 | 0.039 | 0.000 | 0.010 | 0.000 | 0.000 | 0.000 | 0.000 | 0.010 | 0.093 | 0.025 | 0.000 | 0.015 | 0.706 |
| P09 | 0.004 | 0.005 | 0.322 | 0.000 | 0.009 | 0.000 | 0.009 | 0.000 | 0.002 | 0.004 | 0.150 | 0.007 | 0.000 | 0.007 | 0.342 |
| P10 | 0.029 | 0.033 | 0.037 | 0.001 | 0.005 | 0.000 | 0.003 | 0.001 | 0.000 | 0.029 | 0.062 | 0.004 | 0.000 | 0.011 | 0.755 |
| P11 | 0.021 | 0.002 | 0.000 | 0.000 | 0.000 | 0.000 | 0.000 | 0.000 | 0.000 | 0.021 | 0.002 | 0.000 | 0.000 | 0.000 | 0.976 |
| P12 | 0.000 | 0.052 | 0.028 | 0.019 | 0.012 | 0.000 | 0.000 | 0.043 | 0.003 | 0.000 | 0.074 | 0.009 | 0.006 | 0.046 | 0.651 |

### Key cellular signalling protein levels change in muscle fibres with mitochondrial dysfunction

In our second IMC experiment, we used a reduced panel of mitochondrial markers (NDUFB8, MTCYB, MTCO1 and VDAC1) combined with 19 key cell signalling markers to explore how these change with OXPHOS deficiency (Figure 2). The signalling targets (Table 2) were selected to reflect a range of key cellular functions including the mtISR, mitophagy, mitochondrial biogenesis, glycolysis and proteostasis.

**Figure 2.**
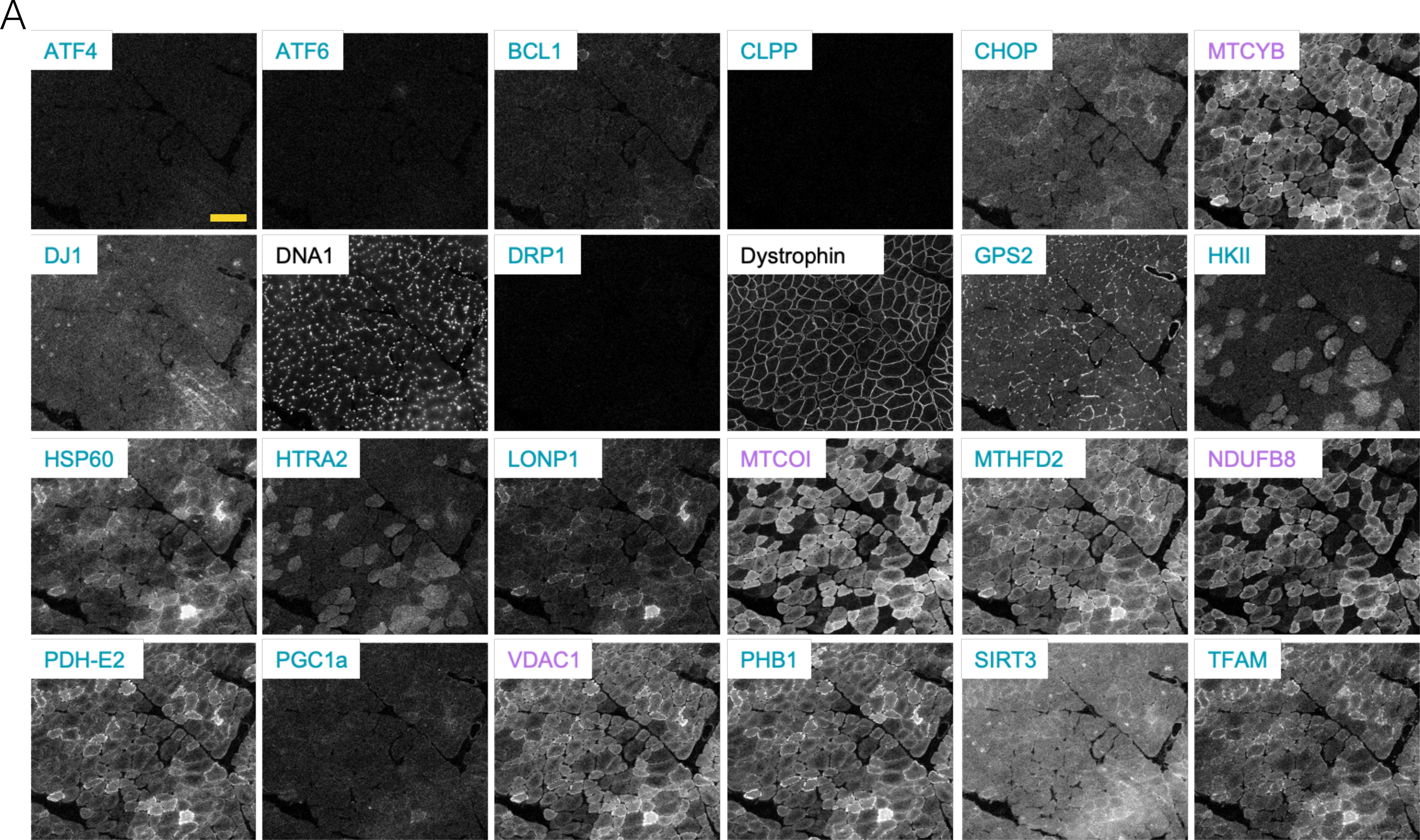
Imaging mass cytometry pseudo-images of mitochondrial and cell signalling targets assessing changes between controls and patients. Pseudo-images showing a range of mitochondrial (purple), cell signalling (turquoise) and cell markers (black) in a muscle tissue sample from a patient.

### Ragged red fibres exhibit a mix of OXPHOS deficiencies

RRFs are an important histological observation in mitochondrial myopathy patients. After identifying RRFs based on morphology and VDAC levels (Figure S1D), we assessed their respiratory chain statuts using our reduced panel of OXPHOS markers to understand if there were particular combinations of OXPHOS deficiency were associated with a RRF phenotype. Interestingly, isolated CIV deficiency was the only class that we could not identify in RRFs. In 7 out of 10 patients with RRFs, the largest group had combined CI, CIII and CIV deficiency (40-100%, n=2-39). In the remaining 3 this was a mix of CI+CIV combined (P02, 51.35% n=19), CIII isolated (P03, 50% n=3) and none (P04, 100% n =1). When looking at the patients combined 60.2% (n=77) of all RRF were deficient in all three complexes (CI, CIII and CIV), 20.3% (n-26) were deficient in CI and CIV, 8.6% (n=11) had no detected deficiency, 6.3% (n=8) had deficiency in CI+CIII, 2.34% (n=3) had isolated CIII deficiency and 0.8% (n=1) isolated CI defciency. It is possible that those with no detected deficiency, had deficiency in CV, but since we do not include a CV marker in our second panel we can not be sure.

### Are there distinct signatures of respiratory chain deficiency?

In order to assess whether cellular response to mitochondrial dysfunction changes dependent on the complexes affected, we used GMM to cluster cells depending on NDUFB8 and VDAC1 for CI deficiency, MTCYB and VDAC1 for CIII deficiency and MTCO1 and VDAC1 for CIV deficiency. GMM was able to separate cells in individual patients into two clusters. We then compared these to the controls to label one cluster as normal or “like controls” and deficient or “not-like controls”. The advantage of this approach is that there is less reliance on the controls as the classes are generated by internal comparison independently of the controls. GMM worked well to identify deficient fibres for CI and CIV but not for CIII, this is because there is also a population of fibres with increased CIII levels and therefore the data cannot be split easily into two. For downstream analysis we required at least three cells per cluster and because CIII deficiency could only satisfy this condition in two patients, we focused our analysis on CI and CIV deficiency (Figure S4).

Having classified each fibre as normal or deficient for CI and CIV, we then grouped them by combination of deficiency into CI+CIV, CI only, CIV only and none. We next sought to compare the different combinations of deficiency. Due to the large variation in the number of fibres that fell into each deficiency group, we performed Bayesian estimation modelling ^26^ to describe the differences in cell signalling targets between muscle fibres from different deficiency groups.

We demonstrated that when compared to cells with no deficiency, CI deficient cells typically have a higher mean levels of PHB1 (9 out of 10 cases effect size (effSz) >0.25), MTCO1 (6 out of 10 cases effSz >0.25), VDAC1 (8 out of 10 cases effSz >0.25), DNA2 (5 out of 10 cases effSz >0.25) and MTHFD2 (8 out of 10 cases effSz >0.25) (Figure 3A). In comparison, CIV deficient cells have a higher mean MTCYB (7 out of 11 cases effSz >0.25) (Figure 3B). Unlike the signalling response observed with CI deficiency where we see a consistent response in all patients, the response to CIV deficiency was much more heterogenous within patients, with P03 and P12 showing a decrease in MTCYB. Similarly, a mixed response is observed for HSP60, TFAM and LONP1.

**Figure 3.**
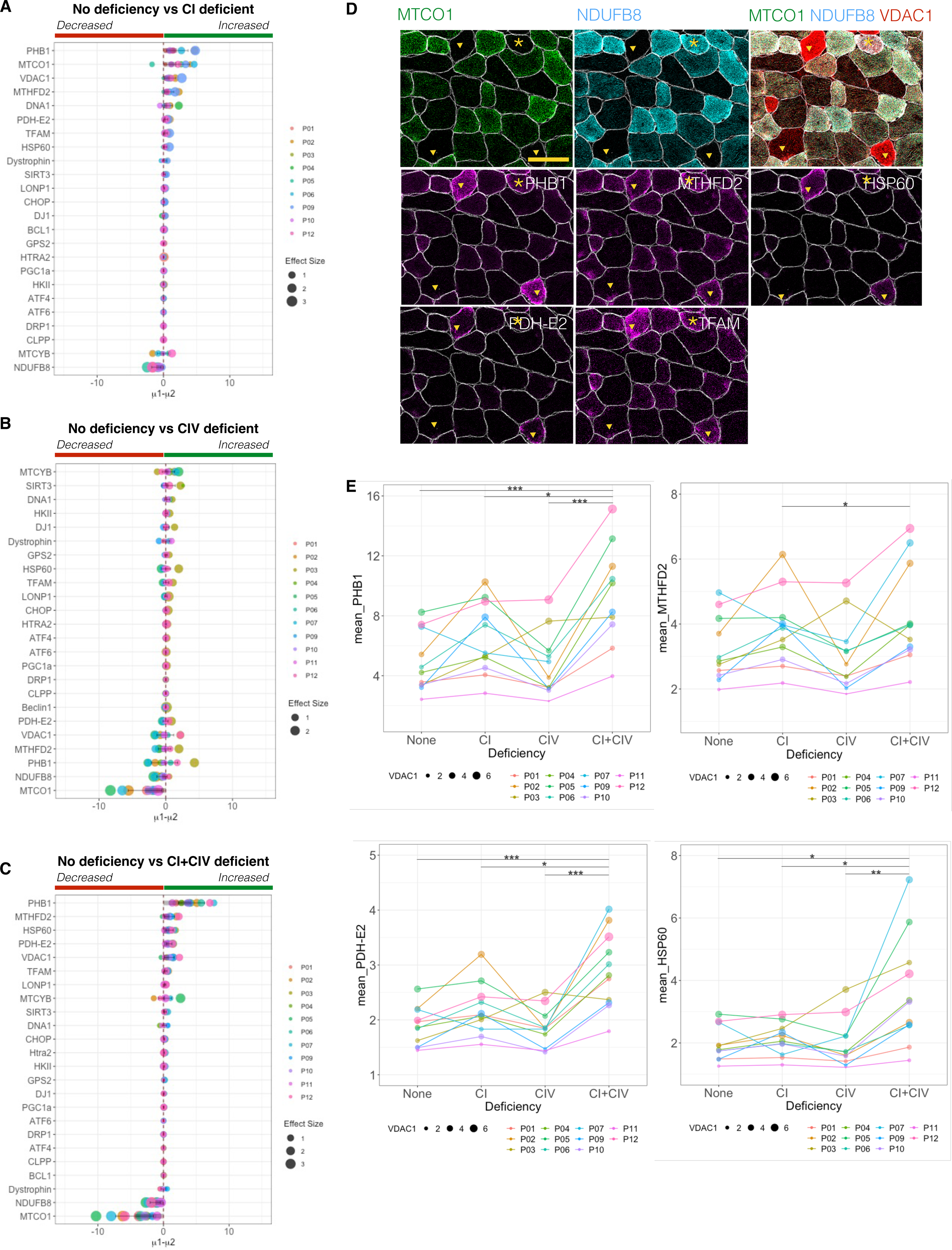
Changes in cell signalling proteins in different classes of mitochondrial oxidative phosphorylation deficiency. Plots show the difference of the mean (µ1-µ2) assessed by Bayesian Estimation versus non-deficient fibres for (A) CI, (B) CIV, and (C) CI+CIV. Proteins are ranked based on the difference of the mean in descending order, with points coloured by patient and point size indicating effect size. (D) Imaging mass cytometry pseudo-images of MTCO1, NDUFB8 and VDAC1, allowing for visual assessment of CI+CIV deficiency alongside PHB1, MTHFD2, HSP60, PDH-E2 and TFAM, and demonstrating the heterogeneity observed in single cell signalling protein levels. Triangles indicate CI+CIV deficient fibres and (*) marking CIV only deficient cells. (E) Plots showing mean intensity levels of PHB1, MTHFD2, PDH-E2 and HSP60 in cells with different combinations of OXPHOS deficiency for each patient. Statistical significance is indicated by a two-tailed Wilcoxon test; *, p<0.05: **, p<0.01 ***, p<0.001.

When compared to normal fibres, CI+CIV deficiency is highly associated with mean increases in PHB1 (11 out of 11 cases effSz >0.25), MTHFD2 (10 out of 11 cases effSz >0.25), HSP60 (11 out of 11 cases effSz >0.25), PDH-E2 (11 out of 11 cases effSz >0.25) and VDAC1 (10 out of 11 cases effSz >0.25) (Figure 3C). Whilst PHB1 and MTHFD2 increases seem to be driven mostly by CI deficiency, increases in HSP60 and PDH-E2 seem to be the result of combined CI and CIV deficiency (Figure 3A and 3C). Looking at the single cell level however, it Is clear to see that mean increases do not necessarily mean that a marker is increased in all cells with CI and CIV deficiency, as can be seen for HSP60 and PDHE2 in the cell indicated by an asterisk (Figure 3D).

To assess whether the relative pattern of changes between the different OXPHOS deficieny groups was homogenous across patients, we plotted mean levels of PHB1, MTHFD2, PDH-E2 and HSP60 (Figure 3E). For PHB1, there was a general trend for the highest levels to be observed in CI+CIV deficient cells with slightly lower levels in CI deficient, followed by CIV deficient and the lowest levels with no deficiency. The one exception was P03, with the lowest levels of PHB1 wth no deficiency and increasingly higher level s with CI deficiency followed by CIV deficiency, and the highest levels in CI+CIV deficiency. A similar pattern with the highest levels in CI+CIV deficiency was observed for MTHFD2, PDH-E2 and HSP60, with the second highest levels in CI deficiency and lowest in no deficiency. The exceptions to this were P03 and P09 for MTHFD2 and PDH-E2 and P03 and P11 for HSP60.

Interestingly while some proteins were found to be elevated with particular combinations of deficiency, the evidence of negative correlations with OXPHOS complexes was less obvious (Figure S5). Negative correlations were observed between the three OXPHOS markers and HK1 and HTRA2 in P01, P03, P05, P06 and P10. P06 and P08 also exhibiting negative correlations between OXPHOS complexes and PDH-E2 and PHB1 and P05 and P06 had negative correlations between NDUFB8 and HSP60 and LONP1.

In accordance with previous observations both in this study and our previous work^20^, OXPHOS markers of complexes which are not deficient in a single muscle fibre seemed to be generally increased, most notably the levels of MTCO1 in the presence of CI deficiency, and MTCYB with isolated CIV deficiency. In comparison, MTCYB seemed on average to be reduced in both CI and CI+CIV deficient cells. Interestingly, an increase in VDAC1, which is indicative of an average increase in mitochondrial mass, is observed in CI and CI+CIV deficient cells, but not in CIV deficient cells.

If we then compare CI and CIV deficiency (Figure S6A), we see a confirmation of the unique signalling signatures for each deficiency group as demonstrated by the comparison with fibres with no deficiency. Comparing isolated CI or CIV deficiency in turn to CI+CIV deficiency, we see that CI+CIV have an increase in average PHB1 levels over and above that observed with CI deficiency alone and that on average the same five top targets distinguish

CI+CIV deficient cells from CI deficient cells or cells with no deficiency (Figure S6B). In general the comparison between CIV and CI+CIV mostly demonstrates the difference in changes between CI and CIV (Figure S6A and C). However, it is important to note that whilst the mean levels discriminate between different populations of deficiency, the distribution of level of some of these proteins varies substantially within each class of deficiency (Figure S6A). Therefore it is also possible that some of these targets are increased with deficiency but not in all fibres and not affecting the mean.

### Is there a cell signalling response associated with the RRFs phenotype?

We then looked to see if there was a difference in the signalling markers between normal and RRF. Interestingly, we find that PHB1 was increased in RRF compared to normal fibres, and is ranked higher in the variable importance in projection plot than VDAC1, the mitochondrial mass marker that we threshold to define RRF (Figure 4). Mean levels of MTHFD2, PDH-E2, HSP60, TFAM and LONP1 were also increased in RRFs (Figure 4). We also found increased levels of DNA1 and DNA2 but when we checked there was no evidence of increased nuclei in RRFs. The signalling pattern observed for RRFs is very similar to that observed in CI+CIV deficient fibres with increased PHB1, MTHFDs, HSP60 and PDH-E2 observed in both.

**Figure 4.**
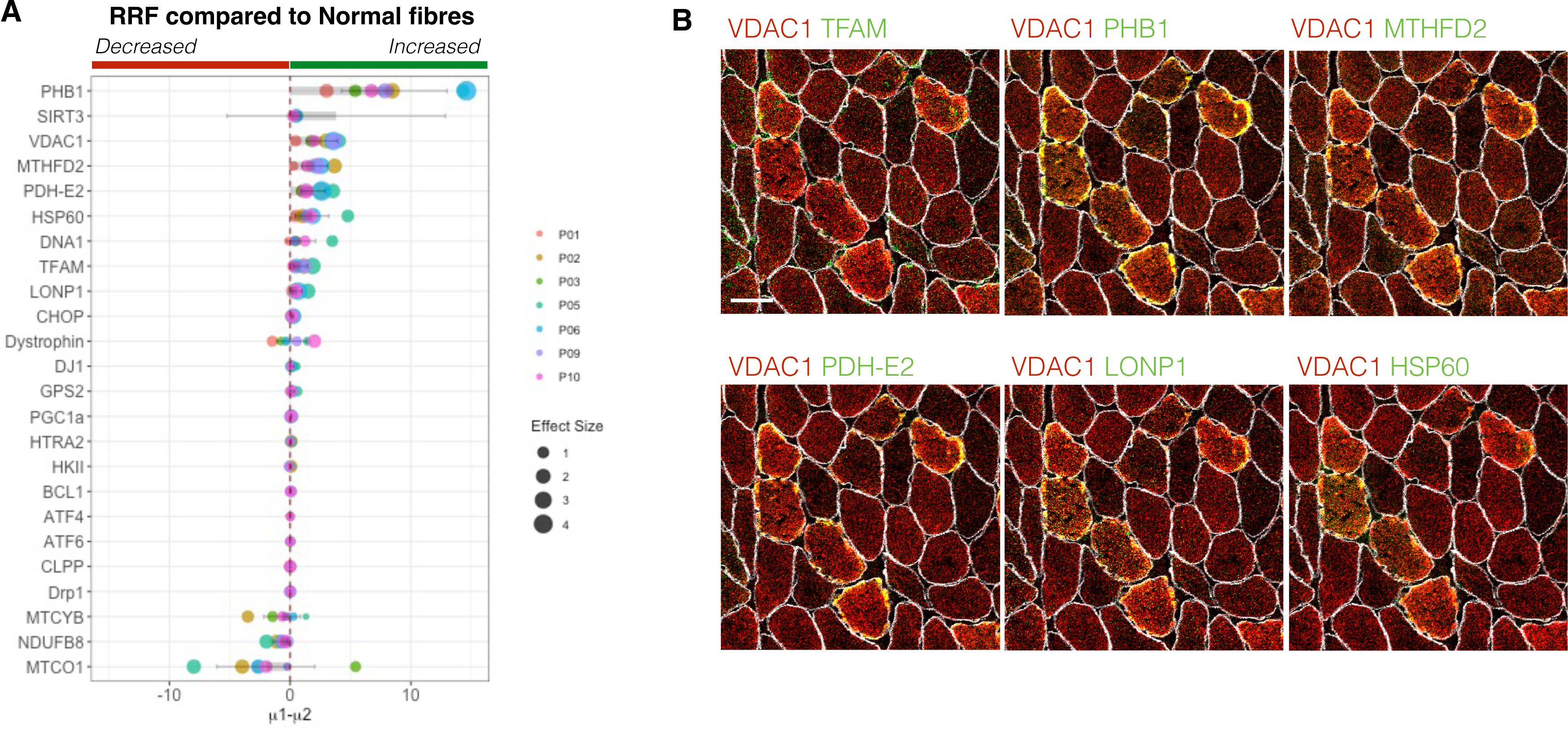
Changes in cell signalling proteins in ragged red fibres (A) Plots show the difference of the mean (µ1-µ2), assessed by Bayesian Estimation, with proteins ranked based on the difference of the mean in descending order. Points are coloured by patient and point size indicates effect size. (B) Merged imaging mass cytometry pseudo-images showing VDAC1 in combination with six signalling markers (TFAM, PHB1, MTHFD2, PDH-E2, LONP1 and HSP60).

Is there a change in nuclear:cytoplasmic ratio of any of the markers with deficiency? Having observed only small changes in proteins known to move between the cytoplasm and nuclear compartments e.g. PGC1α and GPS2, we next sought to see if there was a change in the nuclear to cytoplasmic ratio for these proteins. We extracted mean intensity for each single pixel within each cell. We could then use a threshold for the DNA1 signal to filter the pixels in each cell into nuclear and cytoplasmic, and this classification was plotted and visually compared to images to check for accuracy in the filtering method. Upon examination the resolution of the IMC images appears insufficient to compare nuclear to cytoplasmic ratios, as such high resolution imaging would be required.

## Discussion

Mitochondrial DNA deletions accumulate within muscle fibres of adult patients with mtDNA maintenance disorders and causeing progressive mitochondrial dysfunction.

However, whether the cellular response to such dysfunction changes as the mitochondrial dysfunction progresses has been less clear. Furthermore, it is unclear whether isolated deficiencies in particular complexes would trigger different cellular responses. Here we use IMC to probe a selection of cell signalling markers involved in the mtISR, mitochondrial biogenesis and mitophagy, among other pathways, to understand how these differ with different combinations of OXPHOS deficiency and in RRFs.

Perhaps the most interesting finding is the high levels of PHB1 observed in CI deficient cells and CI and CIV deficient cells, when compared to normal fibres from the same patients. Even more interesting is that PHB1 is not elevated with isolated CIV deficiency. The increase in PHB1 is specific to patients when compared to controls and so has the potential to be a new biomarker, but further investigation is needed and its specificity would need to be confirmed.

Notably the prohibitin complex has been found to be involved in the assembly and degradation of CI in mammals ^27^ and overexpression of PHB1 has been reported to increase CI function, which may explain its selective increase in CI deficient fibres but not isolated CIV deficient fibres. Aside from this, the prohibitin complex has been reported to have numerous cellular functions including a role in the modulation of OXPHOS actibity through interaction with some of the complexes and in the absence of PHB1 instability of mtDNA encoded mitochondrial OXPHOS subunits^28^ . An increase in PHB was previously reported in COX- fibres compared to COX+ in human muscle^16^.

Similarly PDH-E2 is also increased in both CI deficient and CI and CIV deficient cells compared to normal cells from the same patients, but not in CIV alone. PDH-E2 is a subunit of the Pyruvate Dehdrogenase complex a key enzyme involved in the conversion of Pyruvate to Acetyle-CoA for the TCA cycle^29^. A deficiency in Complex I but not Complex IV would shift the NAD^+^/NADH ratio towards NADH as complex I oxidises NADH to NAD^+^. An increase in the levels of PDH-E2 concievably would lead to an increase in substrates into the OXPHOS system. Although this would lead to a viscious cycle for the NAD+/NADH ratio unless a shift to glycolysis is initiated. Murgia et al. found a PDH cluster to be increased in COX- fibres compared to COX+, this fits with our data since tehir proteomics demonstrates these fibres were also CI deficient.

The oxidation of NADH by complex I but not complex IV may also explain why SIRT3, a NAD+ dependent deacetylase, is increased only in CIV deficient cells. SIRT3 is known to regulate energy metabolism and has been shown to regulate glycolysis^30^.However, it should be noted that compared to some of the changes observed in CI and CI+CIV deficient cells the change in SIRT3 in CIV deficient cells is less consistent.

MTHFD2 was also increased in CI and CI+IV deficient fibres in all but one patient. MTHFD2 is an NAD dependent enzyme that is central to the folate-mediated one-carbon metabolism, and it has previously been shown to be increased in the Deletor mouse ^31^ model which carries a TWNK mutation leading to multiple mtDNA deletions in the mouse muscle ^32^. This was later demonstrated to be caused by the disruption of dNTP pools and remodelling of one-carbon metabolism ^33^ and to be regulated by ATF5 in humans ^11^.

TFAM is increased with CI or CI+CIV deficiencies. Since TFAM has previously been shown to increase with mtDNA copy number but is also a regulator of mitochondrial biogenesis, this perhaps suggests that these are mostly triggered by CI deficiency rather than CIV. We previously reported higher levels of TFAM in focal regions of deficiency as well as in some CIV deficient cells ^17^.

Overall, our paper reports key signalling changes in association with mitochondrial OXPHOS deficiency. Our PHB1 findings are striking and warrant future work to understand if PhB1 could be a potential biomarker or therapeutic target in mitochondrial myopathies. Further work will be required to explore some of the interesting targets at higher resolution and to understand whether these changes are specific to mtDNA maintenance disorders, to mitochondrial myopathy, or more generally to OXPHOS deficiency in skeletal muscle more generally.

## Author contributions

Conceived of the idea and planned experiments: AEV Methods development: AEV, CC, TL, KP, AKR, Experimental work and data collection: AEV, DM, AFu. Provision of resources; RC, MM, AFi. Statistical analyses; AEV, CL and CC. Clinical data and tissue: DL, SZ and MD. Data interpretation: AEV, CC, TG, AS, AKR and CL. Drafted manuscript AEV and CC. All authors provided critical revision of the manuscript.

## Acknowledgements

This work was funded by a Henry Wellcome Postdoctoral Fellowship to AEV (215888/Z/19/Z) and the NIHR Newcastle Biomedical Research Centre awarded to the Newacstle upon Tyne Hispotals NHS Foundation Trust and Newcastle University. CC was supported by MJFF grant (grant 15707) and PDUK grant (G-2003) awarded to AKR. We would like to aknowledge the staff of the Newcastle University Flow Cytometry Core Facility for their support and technical expertise, Prof Anu Suomalainen advice and helpful comments, Dr Valentina Perissi for the GPS2 antibody and Dr Thomas Nicholls and Prof Robert Lightowlers for insightful comments.

## Supplementary Figure Legends

Figure S1. Defining respiratory chain deficient cells and Ragged-Red Fibres (RRFs). Gaussian Mixture modelling to define (A) complex I (CI) deficiency, based on NDUFB8 and VDAC1, (B) complex IV (CIV) deficiency, based on MTCO1 and VDAC1, and (C) CIII deficiency, based on CYB and VDAC1. Each point is a muscle fibres and point are colour-coded according to classification in “like control”, in blue, or “not-like control” in red. Black contours show the 0.95 and 0.5 percentiles of the control data. (D) Example of imaging mass cytometry pseudo-image used to identify RRFs based on VDAC1 levels and fibre morphology. RRFs are indicated by red asterisks.

Figure S2. Examples of PlotIMC analysis methods. (A) 2D mitoplots showing respiratory chain levels against VDAC1 levels. Control points are shown in grey and patient points are coloured by respiratory chain protein level with blue being high and red low. Solid grey lines indicate the regression of the control data and dashed lines the 95% predictive intervals of the control data regression model. Above the plot the numbers of fibres above and below the predictive intervals are reported. (B) Stripchart showing mean intensity for each protein. Control cells are shown in grey and patient cells are colour-coded by NDUFB8 (complex I), with blue as high and red as low. (C) Stripchart showing the angle of each cell (point) between the x axis and the regression line with the origin or theta. Points are coloured grey for control and by NDUFB8 for patient cells, with blue as high and red as low.

Figure S3. Heatmap of correlation between mitochondrial markers. Heatmaps show Pearson’s correlation for mitochondrial proteins in each control and patient. Pearson’s correlation is indicated by a colour gradient from 1 (red) through 0 (white) to -1 (dark blue).

Figure S4. Gaussian mixed modelling used to define cells that are complex I and complex IV (CI+CIV) deficient for each patient. Control-like cells are shown in blue and deficient cells in red. Black contours show the 0.95 and 0.5 percentiles of the control data. The percentage of total patient fibres classified as positive or deficient is indicated above each plot.

Figure S5. Heatmap of correlation between mitochondrial markers, cell signalling proteins and cell markers. Heatmaps show Pearson’s correlation for mitochondrial markers, cell signalling proteins and cell markers in each control and patient. Pearson’s correlation is indicated by a colour gradient from 1 (red), through 0 (white), to -1 (dark blue).

Figure S6. Bayesian estimation to compare groups of respiratory chain deficiency. Plots show the difference of the mean (µ1-µ2), assessed by Bayesian Estimation, to compare protein levels in cells with (A) isolated CI vs CIV deficiency, (B) isolated CI vs combined CI+CIV deficiency, and (C) isolated CIV vs combined CI+CIV deficiency. All proteins are ranked in descending order based on the difference of the mean. Point size represents effect size and colour patient ID.

