## Supplementary figures and images for "A stagewise response to mitochondrial dysfunction in mitochondrial DNA maintenance disorders"

### Supplementary Figures 1-6

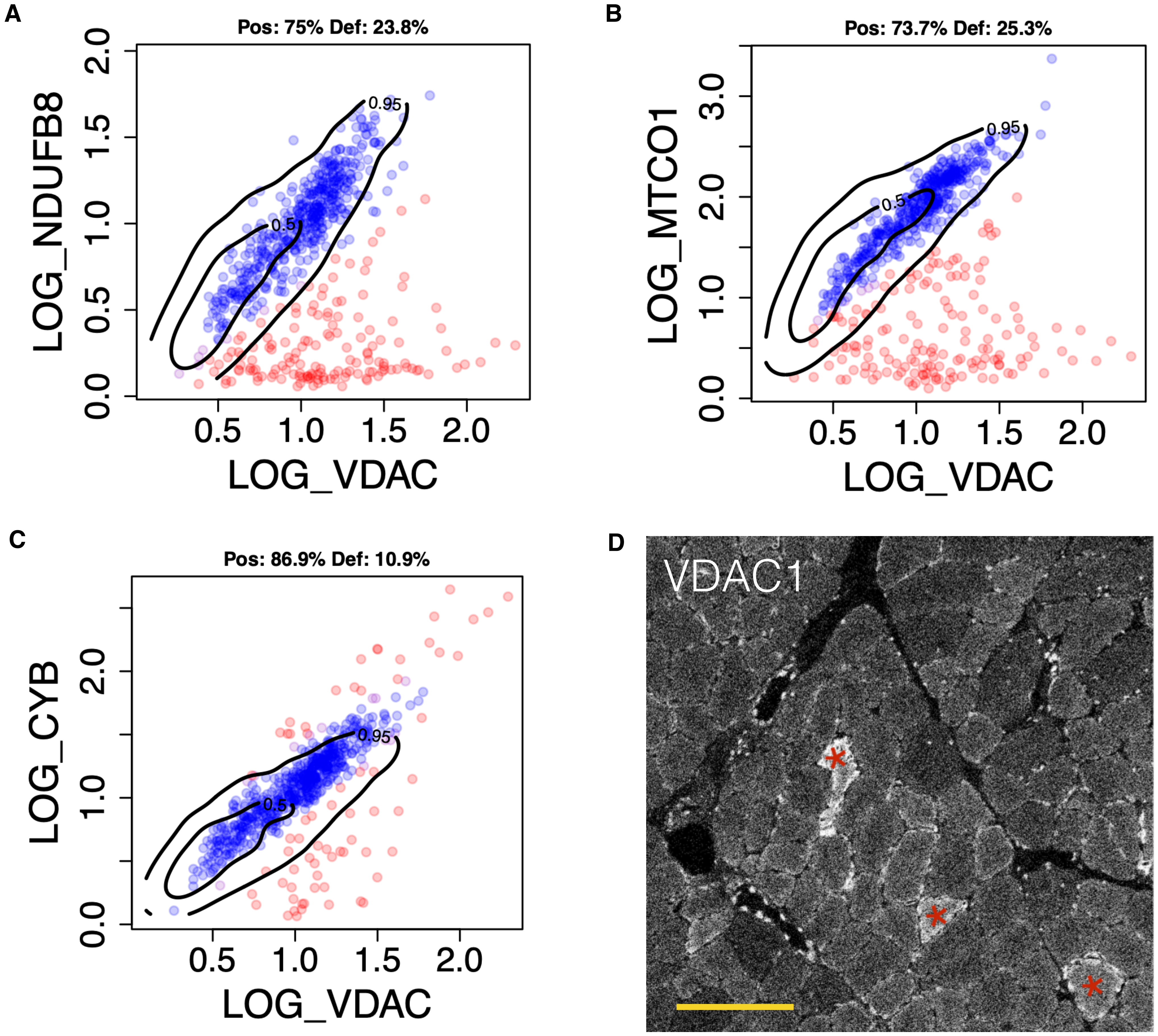

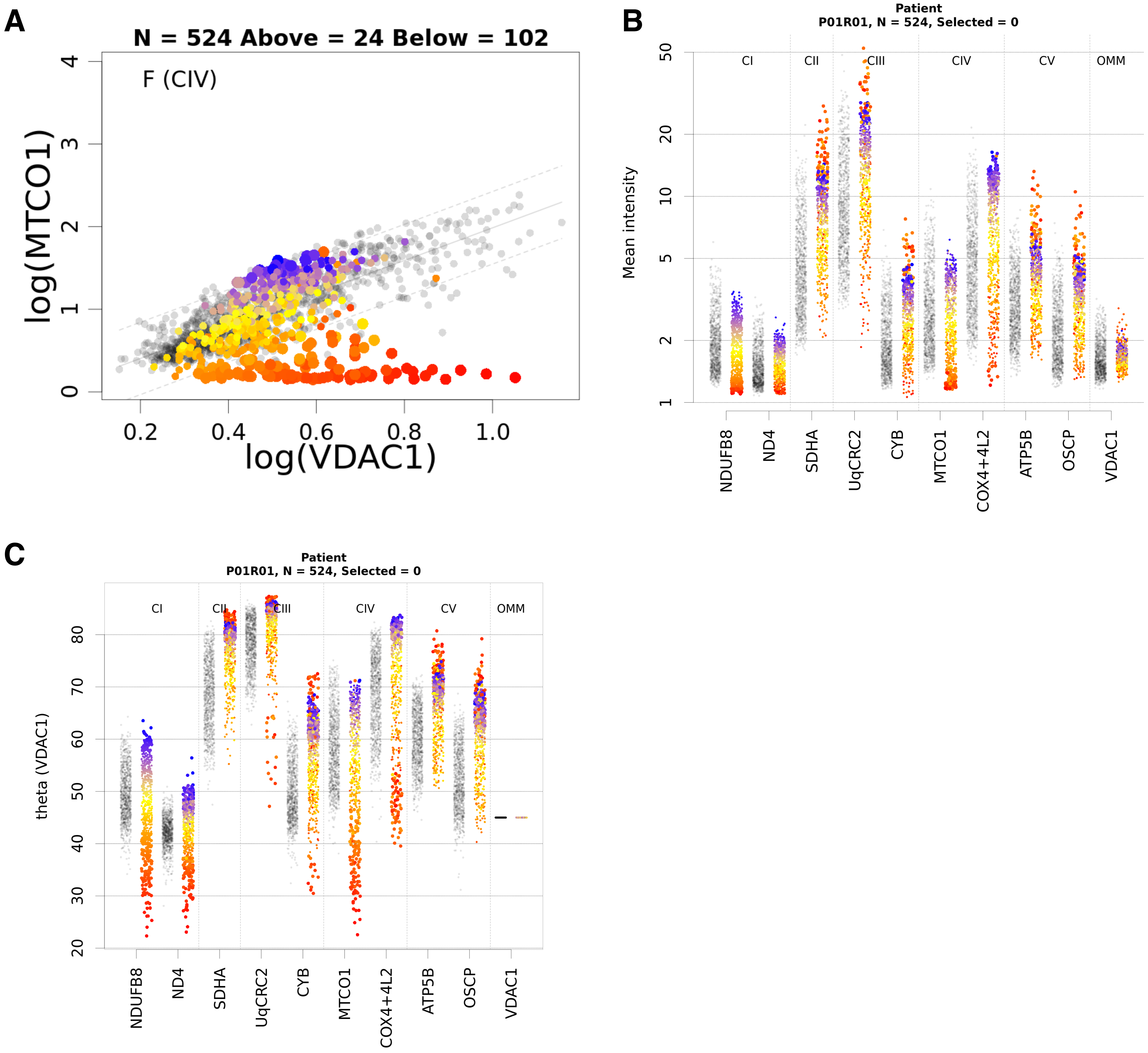

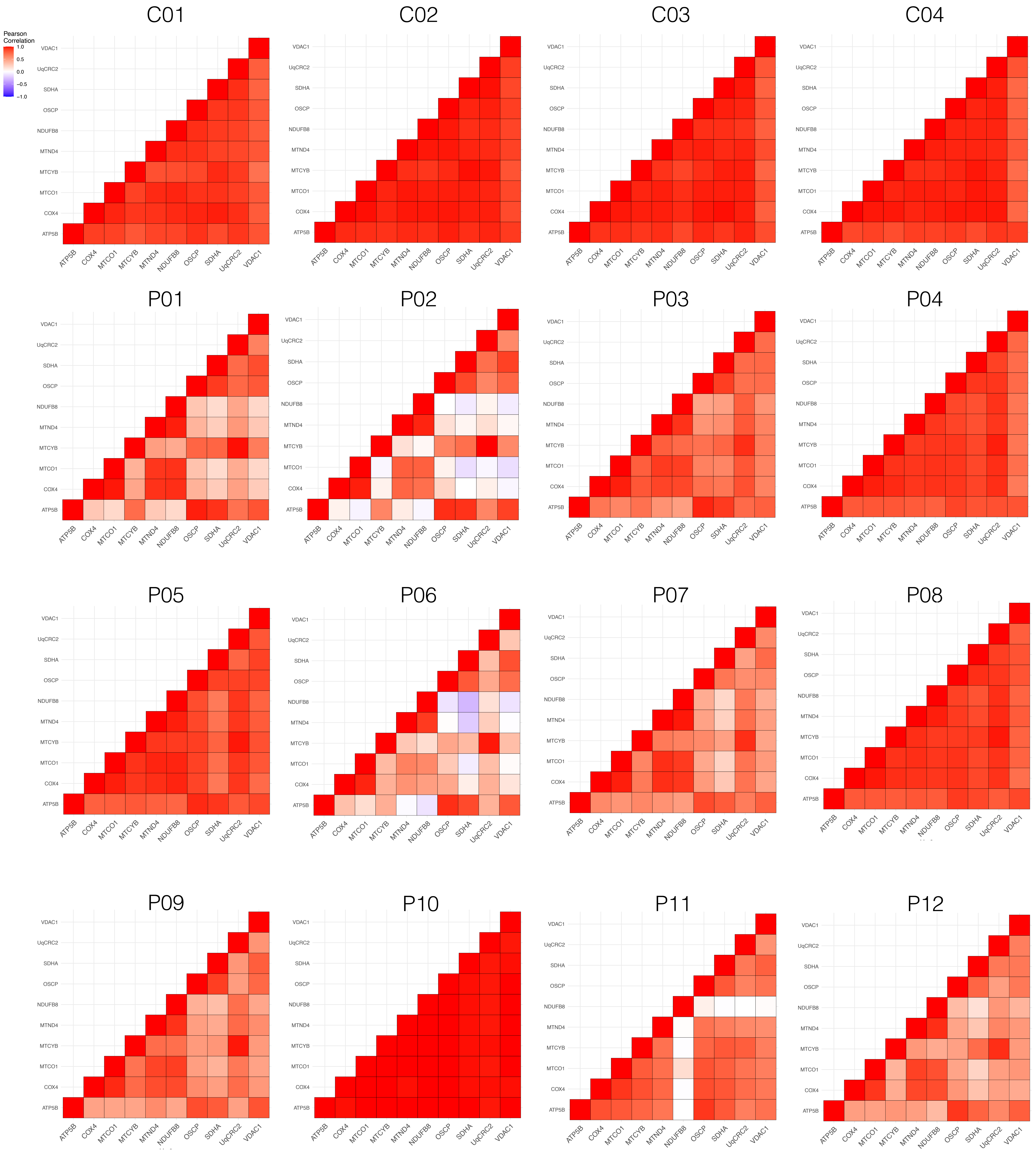

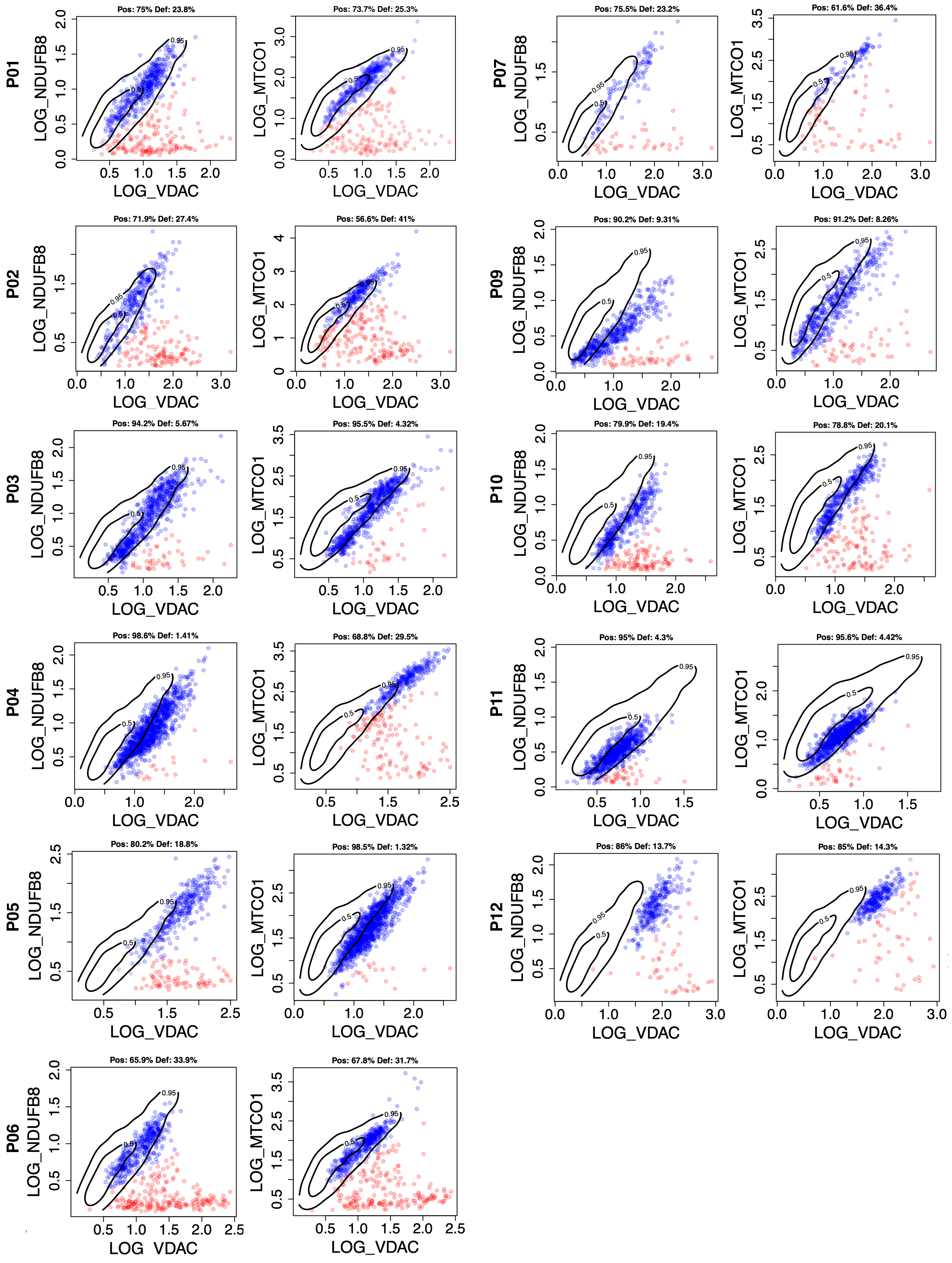

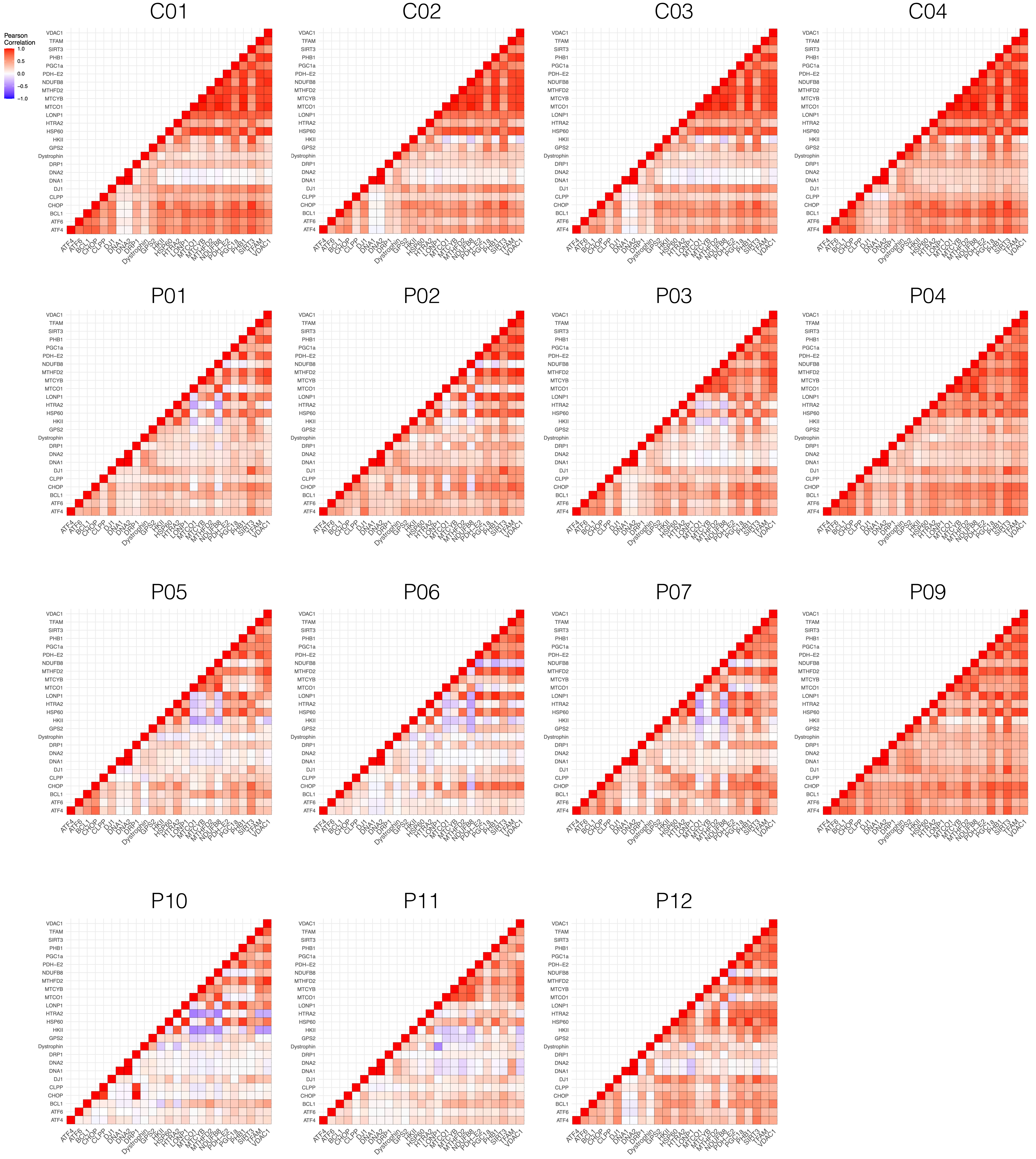

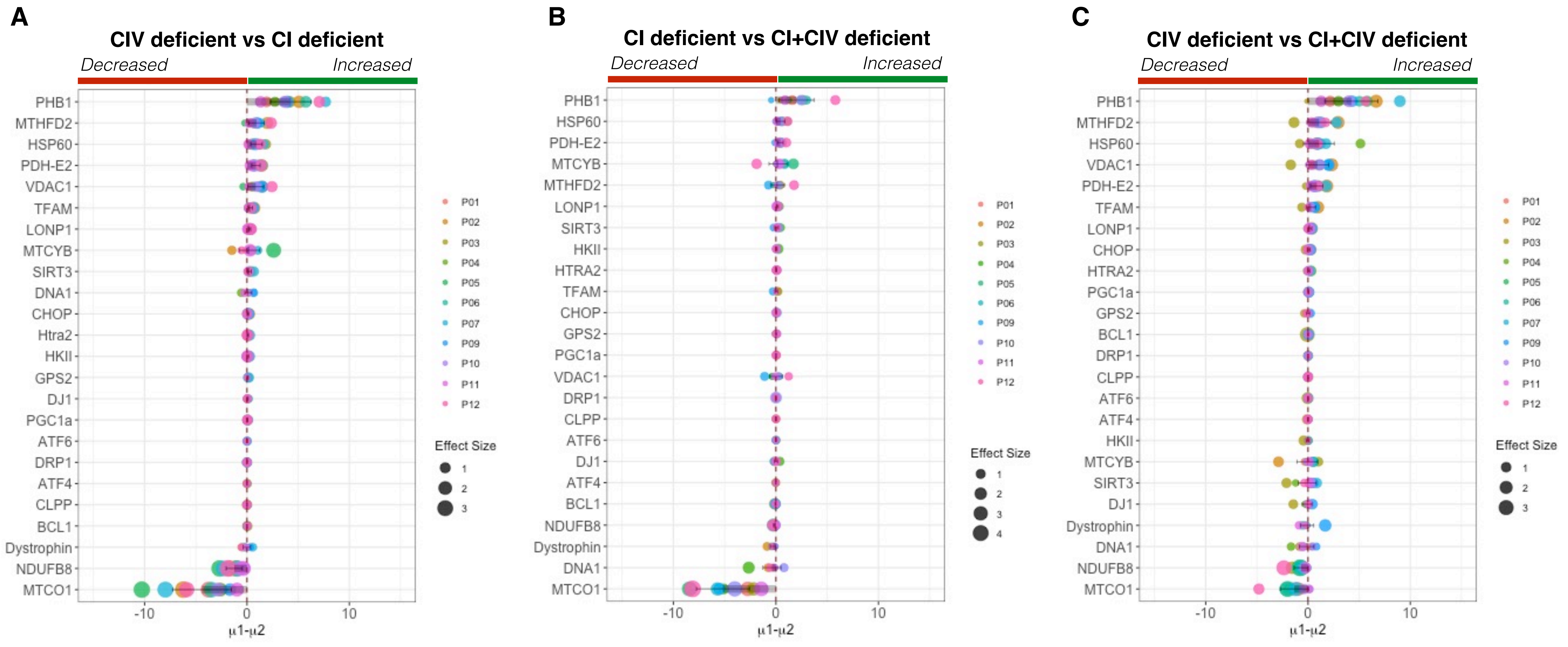
